# Enhancing marine resilience in temperate coastal systems requires the implementation of multiple management strategies

**DOI:** 10.1101/2024.01.13.575496

**Authors:** Jose A. Sanabria-Fernandez, Natali Lazzari

## Abstract

Resilience is a critical property in the current Anthropocene era responsible for keeping temperate coastal systems healthy and preventing their collapse. The cumulative effect of unsustainable practices, however, leads to pollution, physical impacts, or overfishing, which threaten the resilience of these coastal systems. This situation, coupled with a lack of studies on resilience in temperate coastal systems, hampers management toward building more resilient systems. To contribute to this knowledge gap, we assessed the resilience of temperate coastal systems in five marine ecoregions, taking into account three dimensions: biological, environmental, and anthropogenic. Our study identified sixteen top resilient areas to be conserved and thirteen bottom resilient areas with the potential for improved management of anthropogenic factors. In a simulated management scenario within bottom resilient areas, our findings demonstrate that although the management potential of anthropogenic factors varies significantly from 76% for anthropogenic pollution to 27.4% for proximity to the nearest city, the strict management of these factors can enhance resilience by up to 27.97%. Therefore, this research not only contributes to advancing our understanding of resilience in temperate coastal areas but also provides valuable insights into the strategic management of temperate coastal systems, offering a promising pathway towards their sustainability.

## 1. Introduction

In an ever-changing world, resilience emerges as the crucial property that safeguards the integrity of natural ecosystems (Walker & Salt, 2012). This is because resilience buffers sudden or unexpected shifts, ensuring that ecosystems can recover from external perturbation and return to a healthy state (Walker et al. 2004; Walker & Salt, 2012). Therefore, it allows natural ecosystems to adapt to changing environmental, anthropogenic, and biological conditions while maintaining their functions, which decreases the risk of catastrophic ecosystem collapse (Holling, 1973; Folke et al., 2010). As resilience plays a vital role in upholding the functions of natural ecosystems and their associated contributions to people (Cardinale et al., 2012; Reyers et al., 2022), its loss would involve significant and negative consequences for ecosystems, livelihoods, and human well-being (Folke et al., 2004; Agardy, 2005).

Unfortunately, the resilience of natural ecosystems is decreasing at an alarming rate in all ecosystems, from terrestrial (Oliver et al., 2015; Bruno et al., 2019; Boulton et al., 2022) to marine (Lotze et al., 2006; Llope et al., 2011). Specifically in marine ecosystems, there are numerous environmental and anthropogenic factors that can negatively affect their resilience (Maynard et al., 2010; Ladd & Collado-Vides, 2013; Gibbs & West, 2019; EEA, 2020). From an environmental standpoint, storms and hurricanes can trigger loss of resilience by physically damaging the biological community, while rising sea surface temperatures, can lead to knock-on effects such as sea-level rise, heatwaves, or ocean acidification (Hoegh-Guldberg et al., 2007; Doney et al., 2012). Meanwhile, anthropogenic factors such as overfishing, pollution, or physical damage, stand as the main drivers of resilience loss (Ling et al., 2009; Piola & Johnston, 2008; Maynard et al., 2015). For instance, overfishing of marine resources leads to important changes in food webs and ecosystem configuration (Estes et al., 1998); pesticides and chemical contaminants dumped in coastal areas impact the health of marine biological communities by increasing the likelihood of diseases (Folke et al., 2004; Piola & Johnston, 2008); and sea-related activities, such as diving or anchoring, physically impact the seafloor and benthic life by dismantling the benthic structure (Gladstone et al., 2013; Siciliano et al., 2016). Although it is true that both environmental and anthropogenic factors can lead to a loss of resilience, in the short and medium term we are only able to influence the anthropogenic; therefore, it is essential to design management strategies aimed at eliminating or minimizing the impact of these anthropogenic factors on marine resilience (Hughes et al., 2003; Mumby & Steneck, 2008; Morecroft et al., 2012).

To date, little progress has been made in guiding management interventions in coastal systems to improve resilience (Steneck et al., 2019). In 2008, Nyström et al., provided the first attempt of merging the resilience theory with practice by providing empirical indicators of key attributes affecting resilience in coral reefs (i.e., biodiversity, spatial heterogeneity, and connectivity). More recently, Mcleod et al. (2019) argued that resilience-based management, an approach focused on prioritizing actions that influence ecosystems functioning, can support natural processes that sustain tropical marine social-ecological resilience, and Steneck et al. (2019) demonstrated that long-term management interventions focused on reducing algal abundances in coral reefs increases the resilience of juvenile and adult corals. However, while each of these examples share a common objective, which is to enhance the resilience of tropical marine communities, the lack of resilience studies in temperate coastal systems can bring about biased management towards actions that are clearly useful for improving resilience in tropical systems but may not be useful for their temperate counterparts. Although some studies in temperate coastal systems conclude that minimizing environmental and anthropogenic disturbances can enhance the resilience of certain algal species (Perkol-Finkel & Airoldi, 2010; Strain et al., 2015), the number of studies investigating ways to operationalize resilience management in temperate coastal systems remains scarce (Sanabria-Fernandez et al., 2019).

To facilitate the resilience management in temperate coastal systems, the first challenge to overcome is gaining a comprehensive understanding of the existing resilience in the area. Knowing the resilience of temperate coastal systems may help identify geographic areas that require urgent intervention. For example, areas with high resilience may build our knowledge of conservation actions to avoid the future impacts of anthropogenic factors, whereas resilience restoration programs may be more adequate in areas with low resilience. The second challenge to overcome is revealing the anthropogenic factors that contribute to the gain and loss of resilience in temperate coastal systems. For instance, Sanabria-Fernandez et al. (2019) found that anthropogenic physical pressures and pollution obtained minimum scores in areas with high resilience. By identifying these factors and understanding their impact on resilience, it becomes possible to effectively manage them and maintain or increase resilience. Regarding this study, we overcame certain challenges and provided a novel approach to operationalize resilience management. We assessed marine resilience in temperate coastal systems based on seventeen factors belonging to three fundamental dimensions of resilience: biological, environmental, and anthropogenic. Then, we examined whether marine resilience was geographically bounded by ecoregions and whether it showed a homogeneous spatial pattern. Afterward, we investigated the management potential of anthropogenic factors, i.e., the extent to which the management of these factors can be improved and provided specific management interventions tailored to each of the studied anthropogenic factors. Our findings identified both top and bottom resilient areas in all the ecoregions, revealing a heterogeneous spatial pattern of marine resilience. Additionally, by simulating a highly restrictive management scenario for bottom resilient areas, we demonstrated that the effective management of anthropogenic factors can significantly enhance resilience, and that the management potential varies for each of these factors. With this study, we shed light on the resilience management of temperate coastal systems, offering a promising pathway towards their sustainability.

## 2. Material and methods

### 2.1. Study area

We carried out this study in the temperate coastal systems of the Iberian Peninsula, North Africa, the Balearic Islands, and the Canary Islands. We surveyed 300 locations within five marine ecoregions of the world (Spalding et al., 2007): South European Atlantic Shelf, Azores Canaries Madeira, Saharan Upwelling, Alboran Sea, and Western Mediterranean, (hereafter, Atlantic, Canaries, Sahara, Alboran, and Mediterranean, respectively; Fig.1). We focused on these temperate coastal systems because although they host a rich biodiversity gradient and are threatened by multiple anthropogenic impacts (Coll et al., 2010; Alcorlo et al., 2023), the resilience of these systems remains little researched (Sanabria-Fernandez et al., 2019). Moreover, these systems hold significant historical and cultural importance for local communities and represent important fishing grounds and diving spots for national and international tourists. They provide a host of contributions to human well-being (Pascual et al., 2017).

**Figure 1.**
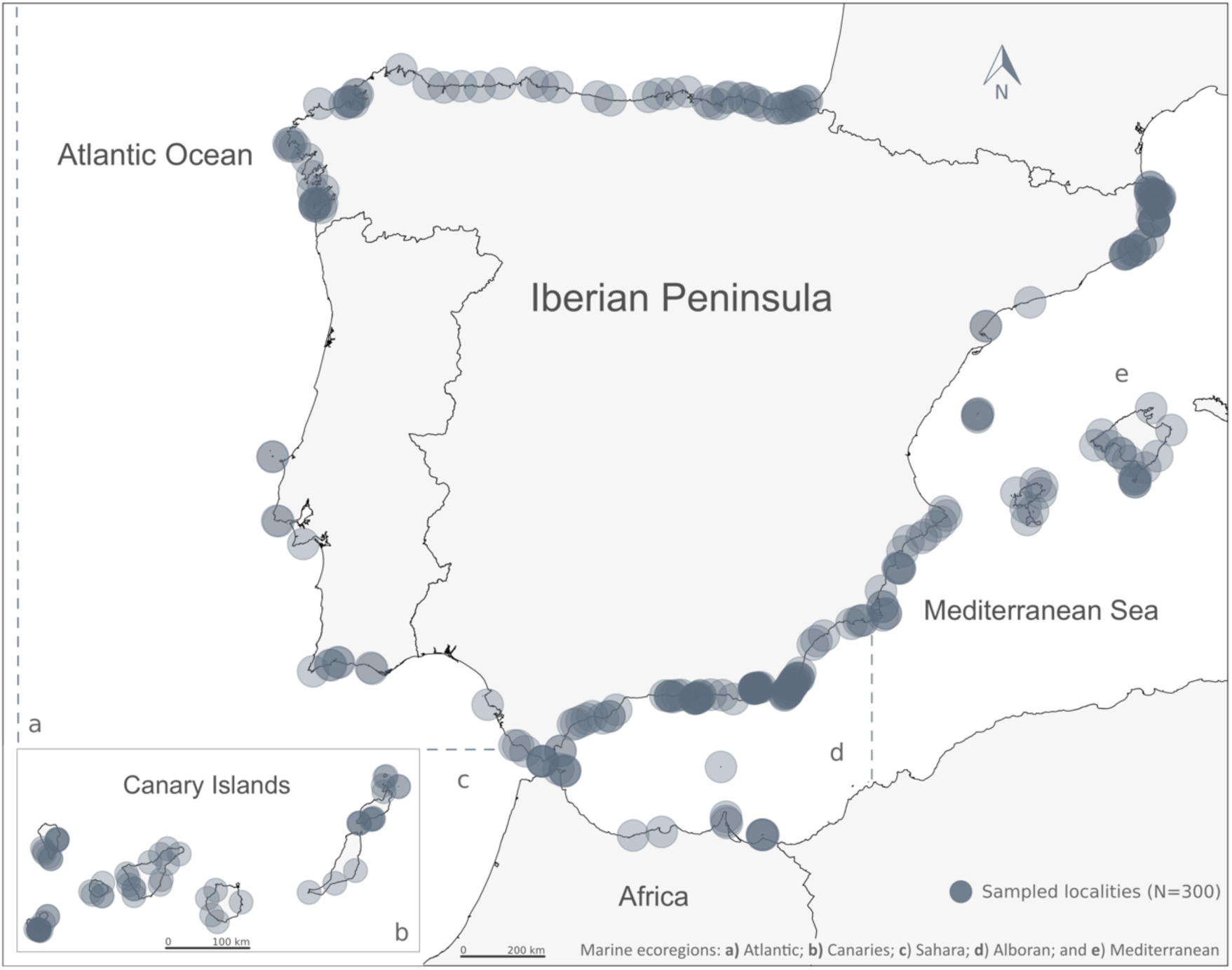
Study area with 300 sampling sites (grey dots; the darkness of shading increases with site overlap). Dashed lines represent the limits of the five marine ecoregions of the world studied: (a) South European Atlantic Shelf, (b) Azores Canaries Madeira, (c) Saharan Upwelling, (d) Alboran Sea, and (e) Western Mediterranean (Spalding et al., 2007).

### 2.2. Factors affecting resilience

Based on previous studies, we defined and quantified seventeen variables as the main factors affecting resilience in temperate coastal systems (Table 1). Factors were classified into biological (nine factors), environmental (three factors), and management (five factors) dimensions following Sanabria-Fernandez et al. (2019) and computed at each surveyed location.

**Table 1.** Seventeen resilience indicators used to quantify the resilience in temperate coastal systems applying the IRIS indicators. For additional details about indicators see Supplementary Table 1.

| **Biological dimension** | **Anthropogenic dimension** | **Environmental dimension** |
| --- | --- | --- |
| **1.** Macroalgal cover | **10.** Fishing pressure | **15.** SSTmax deviation |
| **2.** Community richness | **11.** Anthropogenic pollution | **16.** Nitrate deviation |
| **3.** Abundance of invertebrate herbivores | **12.** Anthropogenic physical pressures | **17.** Phosphate deviation |
| **4.** Abundance of sea urchin fish predators | **13.** Human population |  |
| **5.** Abundance of top predators | **14.** Proximity to the nearest city |  |
| **6.** Fish functional diversity |  |  |
| **7.** Trophic redundancy |  |  |
| **8.** Vulnerability of the fish community |  |  |
| **9.** Abundance of sea urchin invertebrate predators |  |  |

#### 2.2.1. Biological factors

We have calculated nine biological factors affecting resilience: macroalgal cover, community richness, abundance of invertebrate herbivores, abundance of sea urchin fish predators, abundance of top predators, fish functional diversity, trophic redundancy, vulnerability of the fish community, and abundance of sea urchin invertebrate predators. (Table 1; Supplementary Table 1). In doing so, we collected fish, invertebrate and macroalgal information from 300 sampling sites scattered along the coastline of the five marine ecoregions studied, following the standard Reef Life Survey underwater visual census protocol (hereafter, RLS; Edgar et al., 2020). Each sampling site was between 5 and 12 meters deep and represented two or more transects parallels to the coastline. Using RLS, we estimated the abundance of each fish species sighted along a 50m long and 10m wide belt transect, the abundance of each mobile invertebrate species sighted along a 50m long and 2m wide belt transect, and the macroalgal cover from 20 seabed photoquadrats taken every 2.5m along a 50m belt transect (Edgar et al., 2020). Then, we used the biological information collected through RLS to compute the nine biological factors specified in Table 1. For further information regarding the computation of the biological factors, please refer to Supplementary Table 1.

#### 2.2.2. Environmental factors

We used the latitude and longitude of the RLS sampling locations to gather information about the three environmental factors affecting resilience. We collected the mean Sea Surface Temperature (°C), nitrate concentration (mmol·m^−3^), and phosphate concentration (mmol·m^−3^), from the Bio-Oracle database (Assis et al., 2018) using the extract function of the raster R package. All variables were gathered within a 5 arc-minute spatial resolution (Table 1; Supplementary Table 1). We computed the deviation of each environmental factor from the ecoregion’s average. To do so, we calculated the difference between the environmental factor value at a sampling location and the average value of that factor within the ecoregion (Sanabria-Fernandez et al., 2019).

#### 2.2.3. Anthropogenic factors

Concurrently, we gathered information relating to five anthropogenic factors that impact the resilience of the temperate coastal systems: fishing pressure, anthropogenic pollution, anthropogenic physical pressures, human population, and proximity to the nearest city. We obtained this information from national and international public databases (Table 1; Supplementary Table 1).

### 2.3. Quantifying resilience

To compare the real resilience with the resilience of the simulated management scenario, we quantified two resilience indicators, namely the IRIS _Real_ and the IRIS _Simulated_, using the Inclusive Resilience Indicator of a Site – IRIS (Sanabria-Fernandez et al., 2019).

#### 2.3.1. Quantifying real resilience – IRIS _Real_

We estimated the real resilience of the 300 locations surveyed for collecting the biological, environmental, and management factors. To do this, we normalized by min-max (between 0 and 1) of each ecoregion the 17 resilience factors, computing their original factor intensity. Then, we quantified the resilience at each ecoregion following the IRIS method developed by Sanabria-Fernandez et al. (2019), as:

Where k refers to each contributing factor, FI is factor intensity, and N is the total number of factors (17 in our study). Finally, we rename it as IRIS _Real._

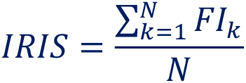

#### 2.3.2. Quantifying simulated resilience – IRIS _Simulated_

To compute simulated resilience, we replaced the original intensity of anthropogenic factors with their maximum management intensity, i.e., 1. This implies minimal pressure from anthropogenic factors due to their management being at its maximum level. Keeping the biological and environmental factors unmodified, we used the formula described in the previous section (2.3.1) for computing the IRIS _Simulated_ at the ecoregion level. The IRIS _Simulated_ represent the maximum hypothetical resilience scores under the lowest anthropogenic pressures, i.e., the most restrictive management scenario.

### 2.4. Unveiling the top and bottom resilience areas

We used the cumulative frequency distribution of the IRIS _Real_ to identify the top and bottom resilient areas. For this, we fitted a dose-response model using a Weibull distribution, which is a non-linear distribution, applying the *drc* R package (Ritz et al., 2015). After that, we detected the lower and upper inflection points (i.e., the thresholds of the distribution) by applying the Unit Invariant Knee point estimation approach, using the *inflection* R package (Christopoulos, 2019). Once we identified the inflection points of the resilience distribution, we classified the IRIS _Real_ scores in bottom, moderate, and top resilient areas. Whereas bottom resilient areas were those with resilience scores between 0 and the lower inflection point, the top resilience areas were those with resilience scores between the upper inflection point and 1. Finally, we classified moderate resilience areas with resilience scores between lower and upper inflection points.

### 2.5. Statistical analyses

To overcome the challenges of managing resilience, we followed three analytical approaches. Firstly, we investigated whether belonging to one ecoregion or another affects marine resilience. To do so, we performed a set of Generalized Additive Mixed Models (GAMMs) with different parameterizations using the *mgcv* R package (Wood, 2017). We fitted the GAMM models with IRIS _Real_ as the response variable and ecoregions as a fixed factor. Secondly, we explored whether the simulated management scenario calculated using the IRIS _Simulated_ significantly enhances the resilience of bottom resilient areas. To do so, we fitted a new set of GAMM models using the IRIS _(Real and Simulated)_ as response variable and two cross fixed factors: IRIS indicators and ecoregions. In both approaches, we included the sampling sites as a random factor to consider the spatial pseudoreplication and the coordinates of the sites as covariates for correcting the spatial autocorrelation (Wood, 2017). We used the minimum AIC - Akaike Information Criteria to select the best-fitted model (Burnham & Anderson, 2002). Thirdly, focusing exclusively on the bottom resilient areas, we assessed whether the management potential, i.e., the extent to which the management of a factor can be improved, varies depending on the factor managed. To that end, we first calculated the management potential as the difference between the maximum score that a simulated anthropogenic factor can achieve, i.e., 1, and the average of 1,000 randomizations of the scores that the real anthropogenic factor has. Afterwards, we fitted a Linear Mixed Model (LME) with the previously calculated management potential as a response variable, the anthropogenic factors, i.e., (AP, FP, APP, HP, and PNC) as a fixed factor, and the sampling sites as a random factor, using lsmeans package (Lenth, 2016). We performed the statistical analyses and graphical representations in R software (R Core Team, 2023).

## 3. Results

### 3.1. Unveiling top and Bottom resilient areas

Resilience was highly heterogeneous across the study area and independent of the marine ecoregion studied (GAMM, F-value=0.88, p=0.48, Fig. 2 and Supplementary Fig. 1). The IRIS _Real_ scores ranged from 0.33 to 0.74, with the Sahara ecoregion showing the lowest and highest resilience scores. We identified 13 bottom and 16 top resilient areas spread across the study area (Fig. 2a & Fig. 2b). We classified the bottom resilient areas as those areas with scores between 0 and 0.4, whereas we classified the top resilient areas as those areas with scores between 0.65 and 1 (Fig. 2a & Fig. 2b). The remaining areas with scores between 0.39 and 0.64, were classified as moderate resilient areas.

**Figure 2.**
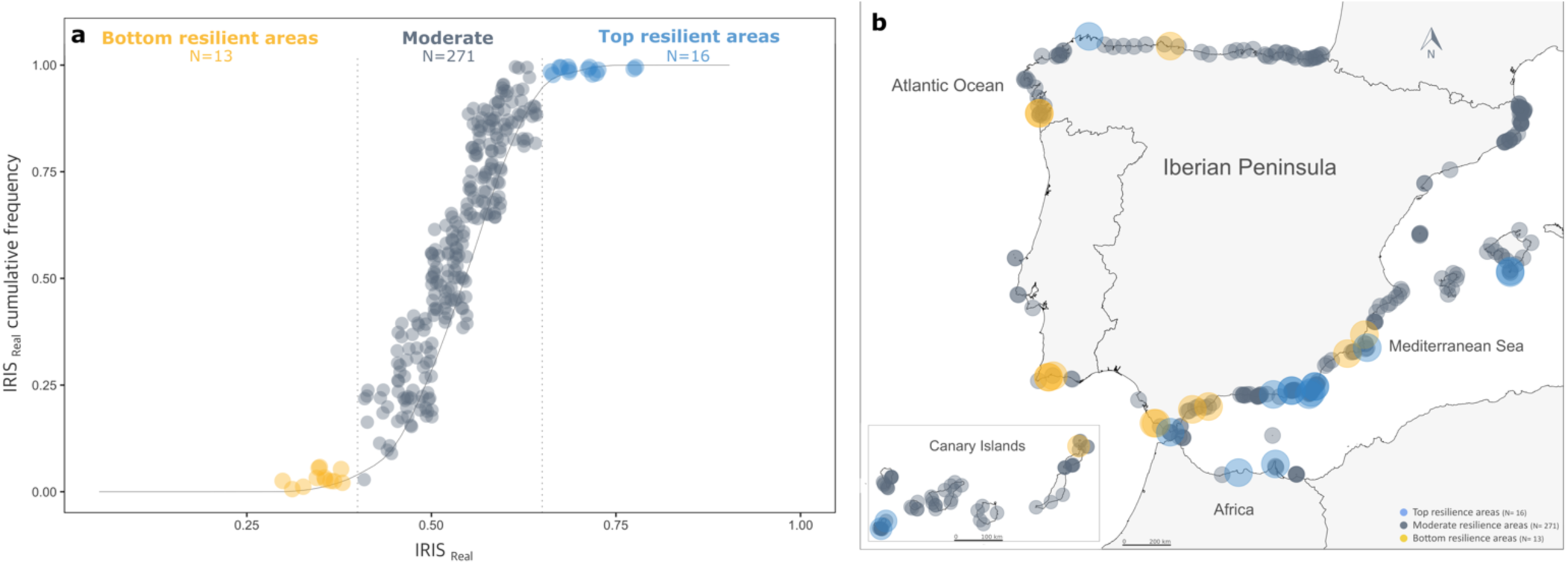
Real resilience of the study site. (a) Cumulative frequency of IRIS _Real_, showing the Weibull distribution (solid line) and the thresholds of the bottom and top resilient areas (dashed lines). (b) Geographical distribution of the bottom, moderate, and top resilient areas.

### 3.2. Enhancing resilience through the management of anthropogenic factors

We found that managing anthropogenic factors could increase resilience by up to 4.79% in the top resilient areas, 11.58% in moderate, and 27.97% in the bottom resilient areas (Table 2, Fig. 3a). On top of that, all anthropogenic factors had a relatively substantial management potential in the bottom resilient areas, which differed according to each factor (LME, F-value= 2580.04, p<0.001, Fig. 3b). This reveals that the management of anthropogenic pollution can be improved by 76%, human population in the coastal area by 56.49%, anthropogenic physical pressures by 44.91%, fishing pressure by 42.81%, and proximity to the nearest city by 27.4% (Fig. 3b).

**Figure 3.**
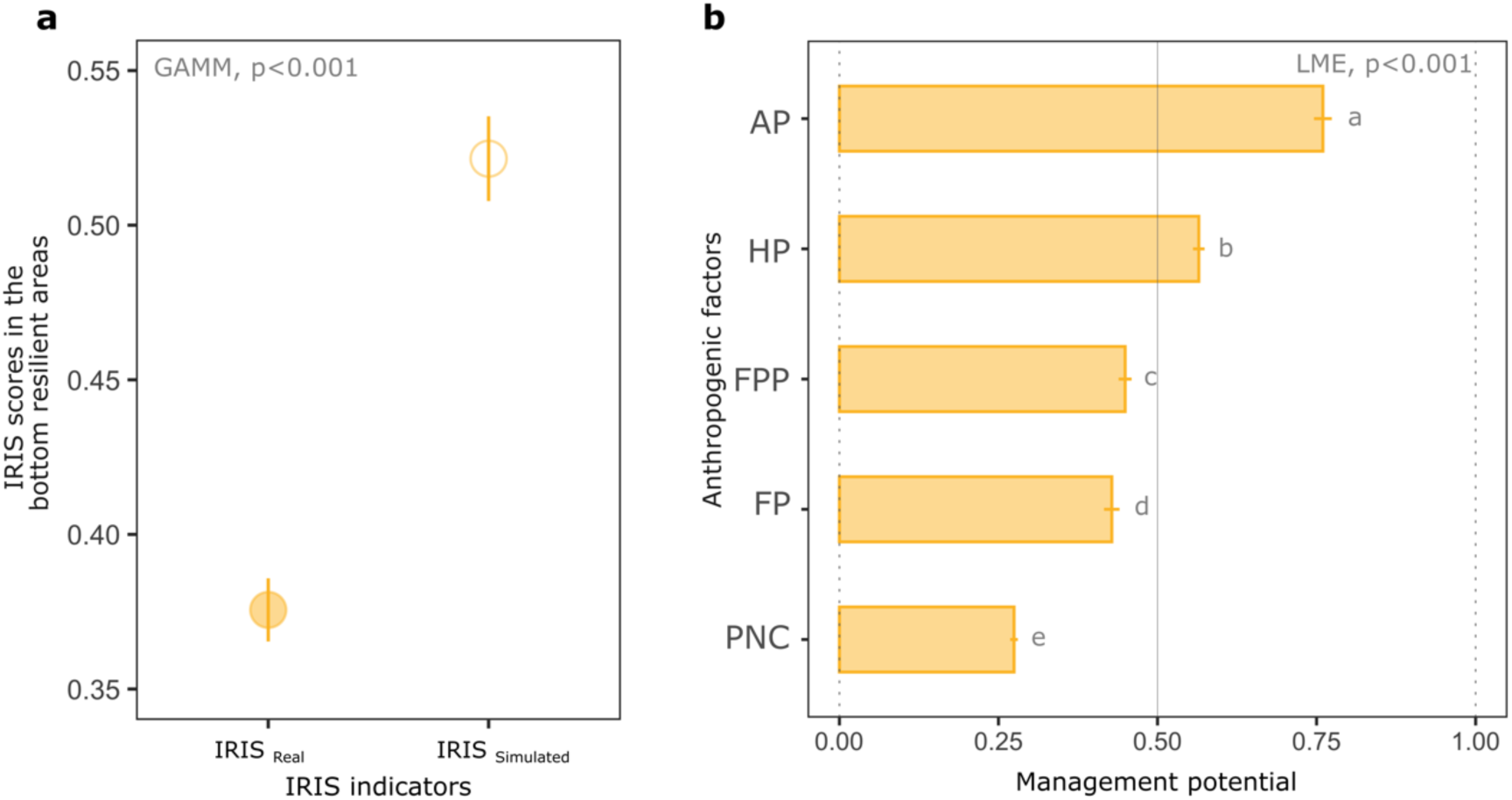
**(a)** Average ± standard error of the IRIS _Real_ and IRIS _Simulated_ in the bottom resilient areas. **(b)** Average ± standard error of the management potential of the anthropogenic factors, namely: AP (anthropogenic pollution); HP (human population); FPP (anthropogenic physical pressures); FP (fishing pressure); and PNC (proximity to the nearest city). Letters indicate significant differences between the investigated anthropogenic factors. The order of the management factors is determined by their management potential.

**Table 2.** Results of the generalized additive mixed models of the bottom resilient areas. We ran the models using two fixed factors, the indicator with two levels (IRIS Real and IRIS Simulated) and the ecoregions with five levels (the ecoregions). Also, we included the sampling sites in the random terms of the model.

| Factors | df | F | p-value |
| --- | --- | --- | --- |
| Ecoregion | 4 | 3.03 | 0.06 |
| Indicator | 1 | 46.8 | <0.001 |
| Indicator*Ecoregion | 4 | 1.77 | 0.19 |

## 4. Discussion

Coastal systems are dealing with the cumulative effect of unsustainable practices such as overfishing, pollution, and physical impacts, among others, leading to an alarming loss of marine resilience (Lotze et al., 2006; Maynard et al., 2015). The combined impact of these unsustainable practices, coupled with a lack of studies on resilience in temperate coastal systems, hampers their effective management toward more resilient systems. In line with this challenge, the present study outlines 16 top resilient areas, i.e., areas with high resilience scores, and 13 bottom resilient areas, i.e., areas with low resilience scores, heterogeneously scattered throughout the studied temperate coastal systems. Furthermore, through simulating a highly restrictive management scenario, we demonstrated that, while each anthropogenic factor possesses its individual management potential, the collective management of these factors can enhance the resilience of bottom resilient areas by up to 27.97%. Hence, our findings advance the understanding of marine resilience, encouraging its conservation and management to build resilient marine temperate coastal systems.

In the top resilient areas identified, there is a narrow margin for improving resilience by managing anthropogenic factors (4.79%). Therefore, management interventions in these areas should focus on increasing their protection status through national, regional, or local institutions with the aim of maintaining their impressive resilience. Similar approaches were taken by Davies et al. (2016) and Green et al. (2009) in tropical systems. They identified priority areas of high resilience in Western Australia and Papua New Guinea, respectively, and designated them as resilient protected areas. These studies are outstanding examples of integrated research on resilience that consider biological, environmental and management factors to reveal resilience patterns and prioritize areas for preservation in the context of climate change. While these examples propose the establishment of marine protected areas as a tool to preserve areas with high resilience, it is important to emphasize that other effective area-based conservation measures, such as in territories managed by indigenous peoples or areas restricted for military purposes, can also be equally effective in conserving and enhancing the resilience of coastal systems (Gurney et al., 2021).

Even though it is crucial to preserve the extraordinary resilience of top resilient areas, we cannot neglect bottom resilient areas. These areas may demand more restrictive and innovative management approaches, yet very restrictive management could enhance resilience by up to 27.97% and anthropogenic factors offer a higher management potential compared to their top resilient counterparts. Therefore, taking action to increase the management of these factors is extremely urgent if we are to prevent further loss of resilience (Maynard et al., 2015; Bruno et al., 2019).

Since our findings show that the management potential of the anthropogenic factors varies for each of them, we were able to create a prioritization list based on this management potential. Anthropogenic pollution was the factor with the highest management potential (76%). Research has shown that the excess of nutrients, pesticides, and other anthropogenic contaminants reduces the resilience of marine communities (Piola & Johnston, 2008; Wernberg et al., 2009). A clear example of this is the ecological collapse experienced by the Mar Menor marine lagoon as a result of the nutrients from the surrounding intensive farming areas continuously entering the lagoon (BCCA, 2020). Reducing the concentration of nutrients, pesticides, and contaminants discharged into coastal areas is the direct action to alleviate this damage. Beyond implementing nutrient discharge restrictions, effective progress can be achieved through the combination of various chemical, physical, and biological techniques in wastewater treatment, including advanced oxidation processes, adsorption, activated sludge, membrane bioreactors, and membrane technologies (Rhind, 2009; Saleh et al., 2020).

Human population, anthropogenic physical pressures, and fishing pressure in coastal areas are the following factors with a high management potential (56.49%, 44.91%, and 42.81% respectively). Population growth and unregulated urban sprawl in coastal areas result in anthropogenic physical impacts on marine biodiversity, which, in turn, affect the resilience of communities (Agardy et al., 2005; Blancas et al., 2010). Limiting or regulating human density across those activities that are most likely to impact marine resilience stands as the best way to manage these anthropogenic factors (Di Franco et al., 2009; Flynn & Forrester, 2019; Giglio et al., 2020). There are already some practical examples of these types of management interventions, such as limiting the number of scuba divers or defining anchoring-free areas, among others. However, these interventions are primarily implemented in areas with a certain level of protection or designed to preserve iconic species such as *Posidonia oceanica* in the Mediterranean Sea (Balaguer et al., 2011). Indeed, the establishment of marine protected areas, specifically no-take zones, both recreational and professional, where anthropogenic activities are completely prohibited has proven to enhance and conserve marine resilience (Edgar et al., 2014; Bates et al., 2014; Perkins et al., 2020). However, solely focusing on restricting anthropogenic activities in protected areas hinders the increase of resilience in those areas lacking protection and that are already highly affected by anthropogenic physical impacts.

Lastly, our results showed that proximity to the nearest city is the anthropogenic factor with the lowest management potential (24.70%). Although the establishment of marine protected areas may also artificially increase the distance between the coastal zone and the nearest city by restricting access, there are other alternative measures that can be adopted to manage this factor. An example of this is the traffic limitations applied in major cities like London, Singapore, or Stockholm to reduce CO_2_ emissions (OECD, 2002; Metz, 2018). Similarly, road closures during the American bison (*Bison bison*) migration season in Wood Buffalo National Park, Canada (https://parks.canada.ca/) and restrictions during sea turtle nesting periods at Mon Repos Beach, Australia (https://parks.des.qld.gov.au/parks/mon-repos), serve as vital measures aimed at safeguarding biodiversity. Restricting the use of those infrastructures that provide access to coastal areas, such as roads and highways, could also artificially increase the distance between the coastal zone and the nearest city, contributing to enhancing the resilience of marine communities in temperate coastal systems.

## 5. Conclusion

Anthropogenic factors stand out as the primary drivers of marine resilience loss in temperate coastal systems. By empirically investigating the marine resilience of the temperate coastal systems of the Iberian Peninsula, North Africa, the Balearic Islands, and the Canary Islands, we identified several management strategies that can be adopted to conserve and enhance the resilience of these systems. Whereas areas with high resilience are the jewels to be conserved to prevent resilience loss, those with low resilience present greater management potential of anthropogenic factors, allowing for a wide range of interventions aimed at restoring and improving their resilience. These management interventions range from the reduction of polluting discharge to the establishment of traffic restrictions, although these interventions should be tailored to the social-ecological peculiarities of the area. Furthermore, we encourage that future studies on temperate coastal systems resilience should address the influence of alternative anthropogenic factors and explore different climate change scenarios. Given that resilience plays an essential role in the health of natural ecosystems and their associated contributions to people, we anticipate that this study will contribute to advancing knowledge for the preservation and enhancement of resilience in temperate coastal systems globally.

## 6. Acknowledgement

We thank all the RLS divers for their invaluable support in data collection during the fieldwork and the projects INBIOMAR and INBIOMAR II, which have been supported by the Biodiversity Foundation of the Ministry for the Ecological Transition and the Demographic Challenge. This article has been drafted and submitted during the Margarita Salas postdoctoral fellowship of JAS-F and NL, funded by the Universitat de Barcelona and the Spanish Ministry of Universities, the Recovery, Transformation and Resilience Plan, and the NextGenerationUE. Finally, we want to acknowledge the Spanish Ministry of Defense for the permits, transportation, and accommodation in the North African military bases.

## Supplementary material

**Supplementary Table 1.**
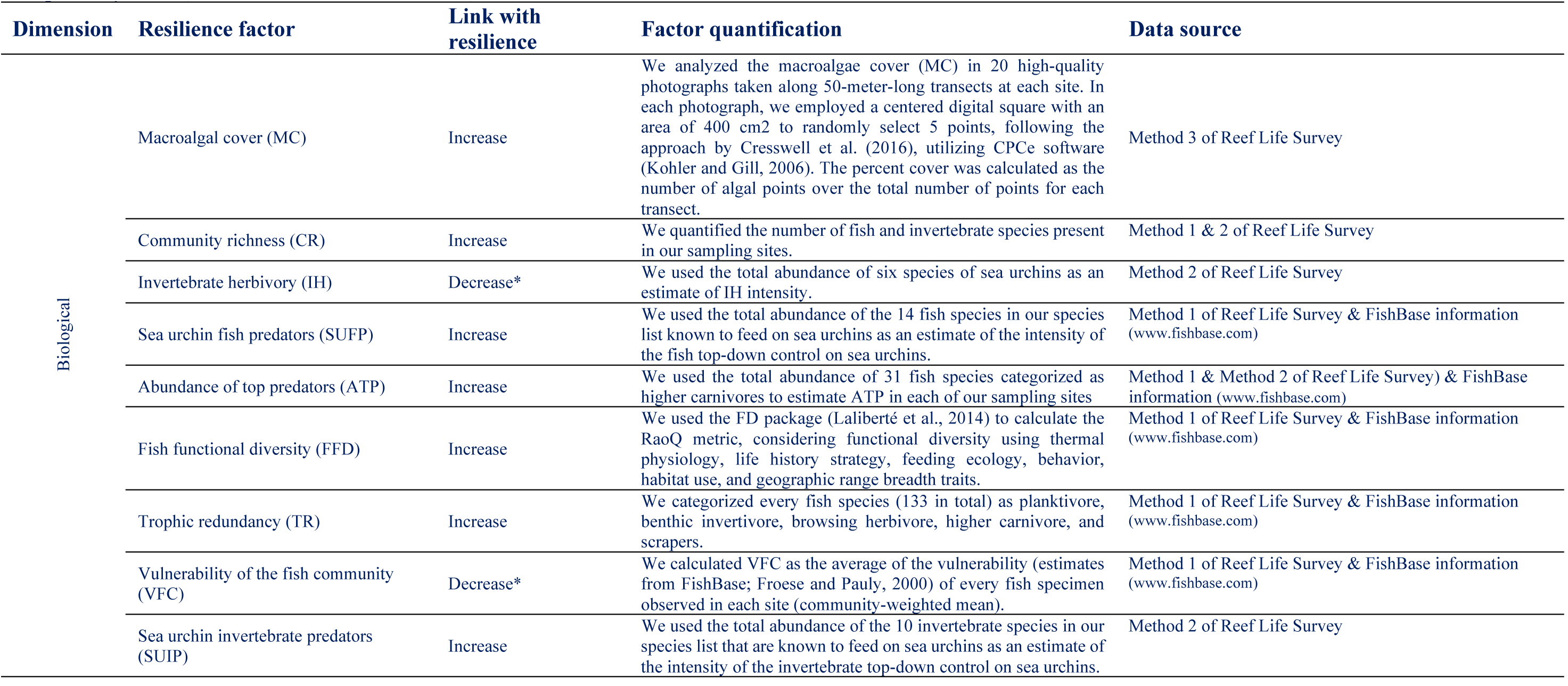
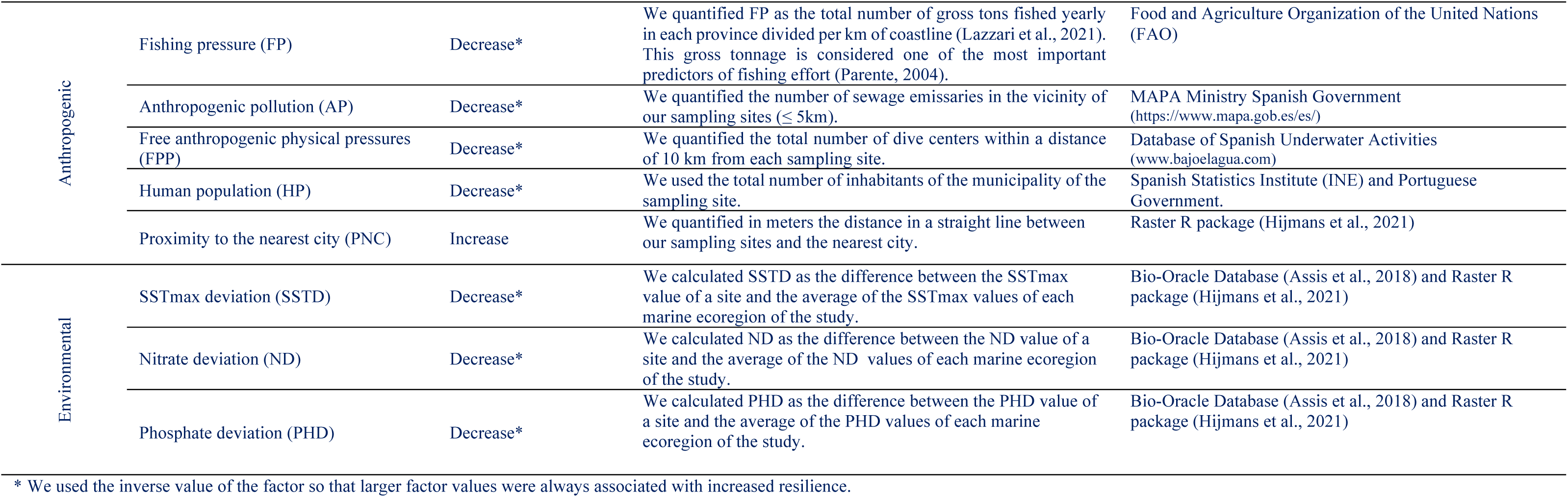
Resilience factors studied to compute IRIS indicators. We selected the factors based on the best available resilience literature in temperate systems (Sanabria-Fernandez et al., 2019).

| Dimension | Resilience factor | Link with resilience | Factor quantification | Data source |
| --- | --- | --- | --- | --- |
| Biological | Macroalgal cover (MC) | Increase | We analyzed the macroalgae cover (MC) in 20 high-quality photographs taken along 50-meter-long transects at each site. In each photograph, we employed a centered digital square with an area of 400 cm<sup>2</sup> to randomly select 5 points, following the approach by Cresswell et al. (2016), utilizing CPCe software (Kohler and Gill, 2006). The percent cover was calculated as the number of algal points over the total number of points for each transect. | Method 3 of Reef Life Survey |
|  | Community richness (CR) | Increase | We quantified the number of fish and invertebrate species present in our sampling sites. | Method 1 & 2 of Reef Life Survey |
|  | Invertebrate herbivory (IH) | Decrease* | We used the total abundance of six species of sea urchins as an estimate of IH intensity. | Method 2 of Reef Life Survey |
|  | Sea urchin fish predators (SUFP) | Increase | We used the total abundance of the 14 fish species in our species list known to feed on sea urchins as an estimate of the intensity of the fish top-down control on sea urchins. | Method 1 of Reef Life Survey & FishBase information (www.fishbase.com) |
|  | Abundance of top predators (ATP) | Increase | We used the total abundance of 31 fish species categorized as higher carnivores to estimate ATP in each of our sampling sites | Method 1 & Method 2 of Reef Life Survey) & FishBase information (www.fishbase.com) |
|  | Fish functional diversity (FFD) | Increase | We used the FD package (Laliberté et al., 2014) to calculate the RaoQ metric, considering functional diversity using thermal physiology, life history strategy, feeding ecology, behavior, habitat use, and geographic range breadth traits. | Method 1 of Reef Life Survey & FishBase information (www.fishbase.com) |
|  | Trophic redundancy (TR) | Increase | We categorized every fish species (133 in total) as planktivore, benthic invertivore, browsing herbivore, higher carnivore, and scrapers. | Method 1 of Reef Life Survey & FishBase information (www.fishbase.com) |
|  | Vulnerability of the fish community (VFC) | Decrease* | We calculated VFC as the average of the vulnerability (estimates from FishBase; Froese and Pauly, 2000) of every fish specimen observed in each site (community-weighted mean). | Method 1 of Reef Life Survey & FishBase information (www.fishbase.com) |
|  | Sea urchin invertebrate predators (SUIP) | Increase | We used the total abundance of the 10 invertebrate species in our species list that are known to feed on sea urchins as an estimate of the intensity of the invertebrate top-down control on sea urchins. | Method 2 of Reef Life Survey |
| Anthropogenic | Fishing pressure (FP) | Decrease* | We quantified FP as the total number of gross tons fished yearly in each province divided per km of coastline (Lazzari et al., 2021). This gross tonnage is considered one of the most important predictors of fishing effort (Parente, 2004). | Food and Agriculture Organization of the United Nations (FAO) |
| | Anthropogenic pollution (AP) | Decrease* | We quantified the number of sewage emissaries in the vicinity of our sampling sites ($\leq 5$km). | MAPA Ministry Spanish Government (https://www.mapa.gob.es/es/) |
|  | Free anthropogenic physical pressures (FPP) | Decrease* | We quantified the total number of dive centers within a distance of 10 km from each sampling site. | Database of Spanish Underwater Activities (www.bajoelagua.com) |
|  | Human population (HP) | Decrease* | We used the total number of inhabitants of the municipality of the sampling site. | Spanish Statistics Institute (INE) and Portuguese Government. |
|  | Proximity to the nearest city (PNC) | Increase | We quantified in meters the distance in a straight line between our sampling sites and the nearest city. | Raster R package (Hijmans et al., 2021) |
| Environmental | SSTmax deviation (SSTD) | Decrease* | We calculated SSTD as the difference between the SSTmax value of a site and the average of the SSTmax values of each marine ecoregion of the study. | Bio-Oracle Database (Assis et al., 2018) and Raster R package (Hijmans et al., 2021) |
|  | Nitrate deviation (ND) | Decrease* | We calculated ND as the difference between the ND value of a site and the average of the ND values of each marine ecoregion of the study. | Bio-Oracle Database (Assis et al., 2018) and Raster R package (Hijmans et al., 2021) |
|  | Phosphate deviation (PHD) | Decrease* | We calculated PHD as the difference between the PHD value of a site and the average of the PHD values of each marine ecoregion of the study. | Bio-Oracle Database (Assis et al., 2018) and Raster R package (Hijmans et al., 2021) |

\* We used the inverse value of the factor so that larger factor values were always associated with increased resilience.

**S. Figure 1.**
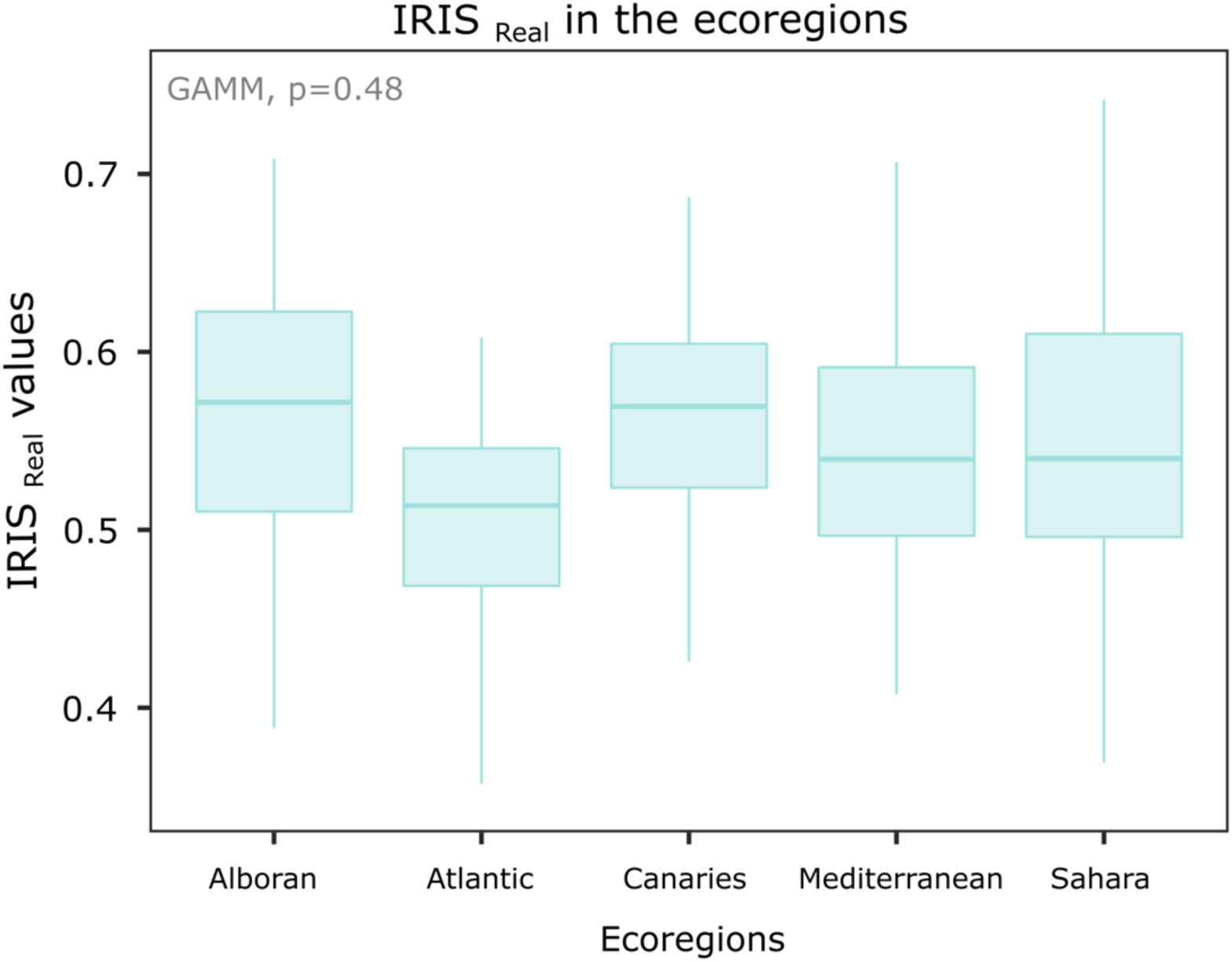
Variation of the IRIS _Real_ values in the five marine ecoregions studied. Boxplots represent the median and 25% - 75% quantiles of the IRIS _Real_ in the ecoregions. The ecoregions are listed in alphabetical order. Alboran: Alboran Sea Ecoregion; Atlantic: South European Atlantic Shelf; Canaries: Azores Canaries Madeira; Mediterranean: Western Mediterranean; Sahara: Saharan Upwelling.

